# A potent agonist-based PROTAC targeting Pregnane X Receptor that delays colon cancer relapse

**DOI:** 10.1101/2024.06.18.599474

**Authors:** Lucile Bansard, Guillaume Laconde, Vanessa Delfosse, Tiphaine Huet, Margaux Ayeul, Emilie Rigal, Quentin Donati, Sabine Gerbal-Chaloin, Martine Daujat-Chavanieu, Baptiste Legrand, Alain Chavanieu, Anthony R. Martin, Julie Pannequin, William Bourguet, Muriel Amblard, Jean Marc Pascussi

**Affiliations:** Institute of Functional Genomics (IGF), University of Montpellier, CNRS, INSERM, Montpellier, France; Institute of Biomolecules Max Mousseron (IBMM), University of Montpellier, CNRS, ENSCM, Montpellier, France; Center for Structural Biology (CBS), INSERM, CNRS, University of Montpellier, Montpellier, France; Institute for Regenerative Medicine and Biotherapy (IRBM), University of Montpellier, INSERM, CHU Montpellier, Montpellier, France

**Author notes:** Contributed equally (Lucile Bansard, Guillaume Laconde, Julie Pannequin, William Bourguet, Muriel Amblard, Jean Marc Pascussi). The authors declare no potential conflicts of interest.

**Keywords:** Colon Cancer, Relapse, PROTAC, PXR

## Abstract

Tumor recurrence is often attributed to drug-tolerant cancer stem cells. We previously demonstrated that down regulation of the Pregnane X Receptor (PXR, NR1I2) decreases chemoresistance of cancer stem cells and prevents colorectal cancer recurrence in xenograft mouse models. These is a lack of PXR antagonists that are appropriate for clinical use. In this study, we report the design and synthesis of a novel PXR agonist-based PROTAC (JMV7048) that induces polyubiquitination and degradation of human PXR protein in an E3 CRBN ubiquitin ligase- and the 26S proteasome-dependent manner. This molecule specifically degrades PXR in colon carcinoma, hepatoma, and pancreatic cancer cell lines, but not in primary cultures of human hepatocytes. Crucially, JMV7048 decreased PXR protein expression in colon cancer stem cells and sensitized them to chemotherapy significantly delaying cancer relapse *in vivo.* PROTACs targeting PXR protein could thus become novel therapeutic agents to enhance cancer cell sensitivity to chemotherapy.

## INTRODUCTION

Colorectal cancer is the third-most diagnosed cancer worldwide, accounting for 10% of cancer incidence and mortality, despite mild advances in early detection and therapy (Siegel *et al*, 2017). Oxaliplatin, irinotecan, and/or 5-fluorouracil are the three major chemotherapeutic drugs used to treat this disease. The addition of targeted drugs to these chemotherapy regimens has further improved survival. Nevertheless, for the vast majority of patients, treatment is unsuccessful once the disease has metastasized to other organs. Therapeutic resistance and tumor recurrence is known to be due to the presence of persister and/or cancer stem cells (Prud’homme, 2012; Adorno-Cruz *et al*, 2015). These cells are highly efficient in metabolizing cytotoxic chemicals, and they are thus only enriched after therapy when their self-renewal capacity kick-starts tumor relapse (Adorno-Cruz *et al*, 2015; Dylla *et al*, 2008). Targeting these resistant cells is therefore crucial for an effective colorectal cancer therapy.

We previously reported that recurrence of colon cancer after chemotherapy treatment is partially due to cancer stem cells overexpressing the Pregnane X Receptor protein (PXR, NR1I2) (Planque *et al*, 2016). PXR is a ligand-activated transcription factor and member of the nuclear hormone receptor superfamily (Lehmann *et al*, 1998). PXR plays a crucial role in the regulation of various metabolic pathways, particularly those involved in the metabolism and clearance of foreign compounds, including drugs, in the liver and intestine (Willson & Kliewer, 2002; Kliewer, 2015). In addition, we and others have shown that PXR participates in the resistance of malignant tumor cells to anti-tumor drugs (Xing *et al*, 2020). For example, PXR expression has been observed and associated with poor survival in various types of cancer, including colorectal (Raynal *et al*, 2010), hepatocellular (Feng *et al*, 2018) and pancreatic cancer (Noll *et al*, 2016). In this context, we showed that the expression of important CSC-related chemoresistance genes (including CYP3A4, ALDH1A1, and ABCG2) and CSC self-renewal depend on PXR expression/activity in colorectal cancer. Additionally, we demonstrated that PXR knockdown, using either RNAi or drug-induced miRNA, delays post-chemotherapy tumor relapse by increasing the chemo-sensitivity of colorectal cancer stem cells (Planque *et al*, 2016; Bansard *et al*, 2022). As a result, after chemotherapy, we (Planque *et al*, 2016) and others (Dong *et al*, 2017) found a negative correlation between PXR expression and recurrence-free survival in colorectal cancer patients. These observations indicate that PXR could be a promising therapeutic target for improving the sensitivity of cancer cells and cancer stem cells to antitumor drugs. The race is now on to translate these recent findings into a clinical therapeutic. However, although nuclear receptors generally respond to a specific set of high-affinity ligands, PXR is activated by a broad spectrum of low-affinity xenobiotics, and crystallographic studies have unveiled a ligand-binding domain (LBD) characterized by a large and conformable binding pocket (Wallace *et al*, 2013; Delfosse *et al*, 2021). This unique feature makes it extremely challenging to identify PXR antagonists through *in silico* or conventional medicinal chemistry approaches (Chai *et al*, 2020). Accordingly, only a few antagonists have been reported to date – SPA-70 (Lin *et al*, 2017), L-sulforaphane (Zhou *et al*, 2007), or ketoconazole (Wang *et al*, 2007). However their efficacy (Poulton *et al*, 2013) and safety make them unsuitable for clinical use (Fuchs *et al*, 2013). Thus, despite the considerable interest in targeting PXR in cancers or in preventing and overcoming drug-induced PXR-mediated drug toxicity and drug resistance, there is currently no PXR antagonist for clinical trials (Kamaraj *et al*, 2022).

Given the challenges associated with inhibiting PXR, we subsequently embraced an alternative strategy. For that purpose, we used an innovative proteolysis targeting chimera (PROTAC) technology to target and degrade PXR. Briefly, this technology has received significant attention as a novel approach for degrading proteins of interest (Paiva & Crews, 2019; Sun *et al*, 2019). PROTACs are bifunctional molecules that hijack the ubiquitin– proteasome system by simultaneously recruiting the target protein and an E3 ubiquitin ligase. This dual engagement triggers ubiquitination and subsequent degradation of the target protein through the proteasome pathway, even for proteins with low ligand-binding affinity. Unlike traditional small-molecule inhibitors, which typically exhibit reversible binding and short-lived effects upon drug removal, PROTAC-induced protein degradation can persist for an extended period (Lai & Crews, 2017). This could be advantageous for achieving prolonged therapeutic responses and potentially lowering the risk of drug resistance (Sincere *et al*, 2023). As a result, PROTACs present an exciting avenue for the development of more effective and durable treatments. Additionally, PROTACs provide a means to interrogate biological processes that cannot be addressed by inhibition alone (Kelly *et al*, 2024). However, designing PROTACs targeting PXR is somewhat challenging due to the complex ligand-binding pocket (LBP) deeply embedded in the PXR LBD (Huang *et al*, 2010). Indeed, the first attempt to create a PXR PROTAC by fusing SPA70 (a PXR inverse agonist (Lin *et al*, 2017)) to thalidomide, a ligand for the E3 substrate receptor cereblon (CRBN), led to the generation of a molecular glue that degrades the GSPT1 translation termination factor instead of PXR (Huber *et al*, 2022). In this study, we present the very first PXR agonist-based PROTAC based on a high affinity homemade human PXR ligand (named hereafter JMV6845 (Benod *et al*, 2008; Lemaire *et al*, 2007)) and thalidomide.

## RESULTS

### Rational development of PXR PROTACs

Given the challenging nature of inhibiting PXR activity, our strategy has been to convert a PXR agonist into a PROTAC to induce the degradation of PXR. Because PXR has the most extensive sequence diversity across vertebrate species in the ligand-binding domain of any nuclear receptor (NR) (Iyer *et al*, 2006), with drastic pharmacologic differences between human and mouse PXRs (Willson & Kliewer, 2002), we selected the potent and human specific PXR agonist JMV6845 previously described (Benod *et al*, 2008; Lemaire *et al*, 2007) (Fig.1A and Fig. S1A). After selecting the PXR ligand, it was crucial to initially identify the suitable position for introducing the linker, a key element between the ligand of the target protein and the ligase, facilitating optimal recruitment of both binding domains. Hence, to elucidate the binding mode of JMV6845, we solved the crystal structure of JMV6845-bound human PXR LBD at a resolution of 2.45 Å (Supplementary table 1). The PXR ligand-binding domain (LBD) adopts the canonical active conformation of nuclear receptors, with its C-terminal activation helix H12 capping the ligand-binding pocket (LBP, Fig. S1B). JMV6845 could be positioned unambiguously in the electron density (Fig.1B) that was well-defined for the entire compound, indicating that JMV6845 is well stabilized in the LBP, in line with its high affinity for PXR. JMV6845 mainly forms van der Waals interactions with LBP residues, and only one hydrogen bond is observed between the nitrogen atom of the sulfonamide moiety and S247 in helix H3 (Fig.1B). Surprisingly, none of the remaining polar atoms of JMV6845 appear to be engaged in a direct or water-mediated hydrogen bond with any LBP residues. The mesityl sulfonamide moiety resides between helices H3 and H11, and it is adjacent to H12, whereas on the other side of the pocket, the benzyl group is nested into the so-called aromatic π-trap composed of F288, W299, and Y306^30^. The central benzimidazole moiety connecting the two aforementioned chemical groups lies between helix H5 and the loop linking helix H2’ and β-strand S1 (loop H2’-S1). Interestingly, despite being fully buried within the LBP, the C-H group of the imidazole ring of JMV6845 (indicated by a red asterisk in Fig.1B) points toward several secondary structural elements that were previously shown to display high intrinsic dynamics (Delfosse *et al*, 2021). Notably, this region of the PXR LBD, comprising helix H2’, the following loop H2’-S1 and H6, is not visible in a number of PXR LBD structures, including when the bulky PXR ligand rifampicin is bound (Chrencik *et al*, 2005) (Fig. S1C). High dynamics of this LBD region could also be observed in the JMV6845-bound PXR structure as the thermal B-factors (representative of the disorder in the crystal) are the highest in the structure (Fig. S1D). We concluded from these observations that the imidazole C-H group could be the most suitable site for the derivatization of JMV6845 in the construction of PXR PROTACs.

**Figure 1:**
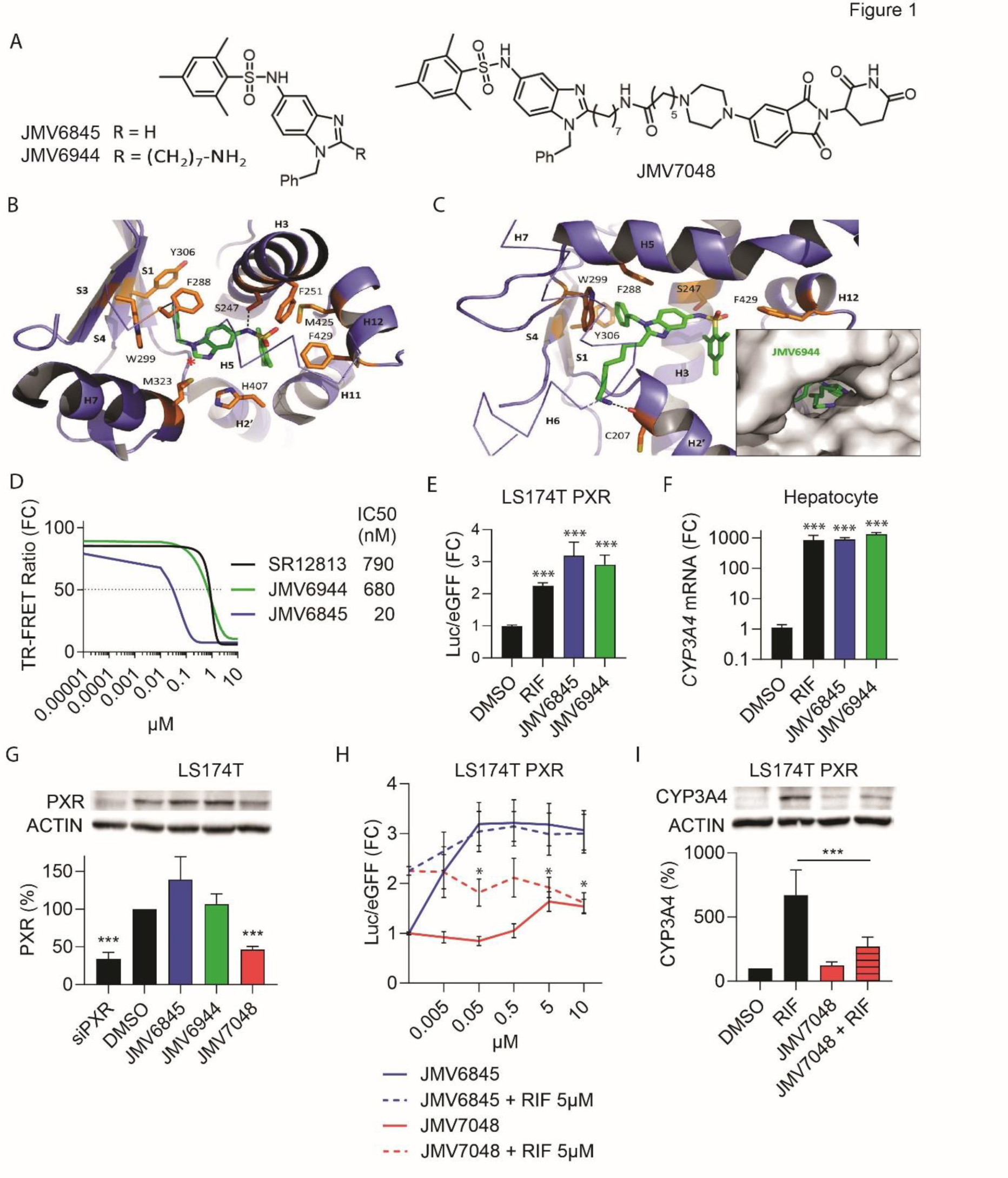
Transformation of a PXR agonist into PXR-PROTAC. **A)** JMV6845, Pre-PROTAC JMV6944 and PROTAC JMV7048 structures. **B)** Close-up view of the interactions between JMV6845 (green) and some residues (orange) of the ligand-binding pocket (LBP) of PXR (Oxygen=red, nitrogen=blue and sulfur=yellow). Key secondary structure elements (α-helices H2’, H3, H7, H11, H12; β-strands S1, S3, S4) are shown and labeled. The red asterisk denotes the C-H group of the imidazole ring serving as a derivatization site of JMV6845. **C)** Close-up view of the interactions between JMV6944 and some residues of the LBP of PXR. The black dashed line denotes a hydrogen bond between the amine moiety of the side chain of JMV6944 and the main chain carbonyl group of cysteine 207. The inset displays a surface representation of PXR-LBD (in grey) showing how the JMV6944 side chain finds its way to the protein surface. **D)** TR-FRET competitive assay between a fluorescent agonist and SR12813 (agonist of PXR) or JMV6845 or JMV6944. TR-FRET ratio (520nm/490nm x 10^4^) are expressed as mean ± SEM (n >3) and normalized to DMSO (%). **E)** PXR transcriptional activity was determined in LS174T cells transfected with a CYP3A4 luciferase reporter plasmids, a CMV-eGFP and a PXR expression vectors. Cells were treated 24 hours with 5μM Rifampicin (RIF, PXR agonist), JMV6845 or JMV6944. Data are expressed as mean ± SEM (n=3) of luciferase/eGFP ratio, normalized to DMSO condition. **F)** RT-qPCR analyses of *CYP3A4* mRNA expression in human hepatocytes treated 24 hours with 5 µM RIF, JMV6845 or JMV6944. Data are expressed as mean ± SEM (n=3) and normalized to DMSO condition. **G)** Western blot analysis of PXR expression in LS174T cells treated 24 hours with 5µM with the indicated compounds. Data are expressed as mean ± SEM (n≥3) of PXR/ACTIN ratio and normalized to DMSO (%). **H)** PXR transcriptional activity was determined in same cells as panel E. Cells were treated 24 hours with an increasing concentration of JMV6845 or JMV7048 and then incubated 24 hours with RIF at 5µM. Data are expressed as mean ± SEM (n=3) of luciferase/eGFP ratio, normalized to DMSO condition. **I)** Western-blot analysis of CYP3A4 expression in LS174T cells over expressing PXR after pre-treatment with or without JMV7048 at 5µM for 24h then co-treated with or without induction by RIF at 5µM for 24h. Data are expressed as mean ± SEM (n=3) of CYP3A4/ACTIN ratio and normalized to DMSO (%). ***, p<0.0005.

Thus, guided by the binding mode of JMV6845 within the PXR LBD, we introduced an aminoheptyl chain onto its imidazole C-H group, resulting in the analog JMV6944, (Fig.1A). As expected, the crystal structure of the JMV6944-bound PXR LBD solved at 2.10 Å resolution (Supplementary table 1 and Fig.S2A and S2B) unequivocally revealed that, owing to a large shift of helix H2’ (Fig.S2C), the aminoheptyl chain weaves its way between helices H2’, H6 and H7 to reach the outer surface of the domain (Fig.1C). The free amine function of the linker forms a hydrogen bond with the surface residue C207, further stabilizing the interaction of JMV6944 with PXR (Fig.1C). Competitive binding assays using time-resolved fluorescence resonance energy transfer between a fluorescent PXR ligand and the purified human PXR LBD (LanthaScreen TR-FRET PXR Competitive Binding Assay) revealed that JMV6944 can bind PXR (IC_50_=680nM) (Fig.1D). Despite having a lower affinity compared to JMV6845, JMV6944 still surpasses the affinities recorded here for compound SR12813 (IC_50_=790nM), known as one of the most potent human PXR agonist (Orans *et al*, 2005). As predicted, both JMV6845 and JMV6944 yielded a robust response in an *in vitro* PXR luciferase reporter assay (Fig.1E) where the CYP3A4 distal and proximal promoters control the luciferase reporter gene (Drocourt *et al*, 2002). In agreement, RT-qPCR analysis confirmed that JMV6944 is a potent inducer of CYP3A4 mRNA expression in freshly isolated human primary human hepatocyte cultures (Fig.1F), considered the gold standard for investigating PXR activity (Pascussi *et al*, 2000). Similar results were observed in LS174T cells (Fig.S3A), known to express functional PXR (Raynal *et al*, 2010; Wang *et al*, 2011). Altogether, these structural and functional data prompted us to further explore the development of PXR degraders using JMV6944, to which a variety of chemical fragments inducing PXR degradation could be easily attached via the amino group of the linker. In light of these results, which demonstrate the effective binding of JMV6944 to the LBD of PXR and the accessibility of the amine function for derivatization, we designed and synthesized JMV6944-based PROTACs.

Our initially focus was on using the cereblon (CRBN) thalidomide ligand, which has led to the development of potent PROTACs targeting various proteins, several of which are currently undergoing clinical trials (Jiang *et al*, 2023). A first series of PROTACs was synthesized by modulating the nature of the linker between the two warheads (Undisclosed data). This investigation enabled us to identify a lead compound, JMV7048, which incorporates the CRBN-targeting compound thalidomide, and JMV6845 targeting PXR, linked by a piperazine hexanamide moiety (Fig.1A). This molecule was synthesized following the procedure outlined in Supplemental Scheme S1 and described in the referenced patent WO2022243365. Subsequently, JMV7048 was evaluated based for its ability to act as PXR degraders, in the LS174T human colon carcinoma cell line. JMV6845 and JMV6944 were also tested as negative controls for PXR degradation. As depicted in Fig.1G, a 50% reduction of PXR protein level was observed after treating LS174T cells with 5µM JMV7048 for 24 hours. As expected, PXR expression remained unchanged upon treatment with JMV6845 or the synthetic intermediate JMV6944 lacking a E3 ligase ligand. Building upon this lead compound, we made slight modifications to the linker by extending its length by one or two methylene units, resulting in JMV7505 and JMV7506, respectively (Fig.S3B). Western blot analysis revealed that the introduction of a single methylene unit within the JMV7048 linker (JMV7505) had no impact on PXR degradation efficiency, while the addition of two units (JMV7506) completely prevented PXR degradation (Fig.S3C). These findings underscore JMV7048 as our lead candidate for PXR degradation. Subsequently, we examined whether JMV7048 also promotes inhibition of PXR signaling by assessing its effect on rifampicin-mediated PXR activation. We performed PXR luciferase reporter assays where cells were first pretreated for 24 hours with increasing concentrations of JMV7048 or the negative control JMV6845, and then treated for 24 hours with 5µM rifampicin (Fig.1H). The results showed that while JMV6845 and rifampicin yielded robust and additive responses, JMV7048 inhibited rifampicin-mediated PXR transactivation in a dose-dependent manner. In agreement, JMV7048 also inhibited rifampicin-mediated endogenous CYP3A4 expression (Fig.1I and Fig.S3D). All these findings suggest that JMV7048 likely acts as a PROTAC by binding to PXR and inducing its degradation in colorectal cancer cells. Consequently, a serie of supplementary tests has been undertaken to unequivocally establish that JMV7048 indeed acts as a PROTAC.

### JMV7048 is a bona fide PXR PROTAC

Because the first attempt to create a PXR PROTAC led to the generation of a molecular glue (SJPYT-195) instead of a bona fide PXR PROTAC^8^, we first performed comparative time-course analyses of PXR protein and mRNA expression upon JMV7048 treatment. As shown in Fig.2A, the kinetic of PXR protein loss induced by JMV7048 was rapid and not related to a decrease of PXR mRNA level (Fig.2A), unlike to the SJPYT-195 molecule^8^, indicating that these effects are independent of transcription effects. In addition, both real-time cell proliferation assays, measured with the xCELLigence RTCA impedance technology, and viability analysis using Sulforhodamine B assay (Fig.2B and Fig.S3E) demonstrate that JMV7048 shows no cytotoxic effects in all tested CRC cell lines. This is evident at a concentration as high as 20μM applied for 72 hours, in comparison to the active metabolite from irinotecan (SN38), which is the main chemotherapy treatment of colorectal cancer patients (Fujita *et al*, 2015).

**Figure 2:**
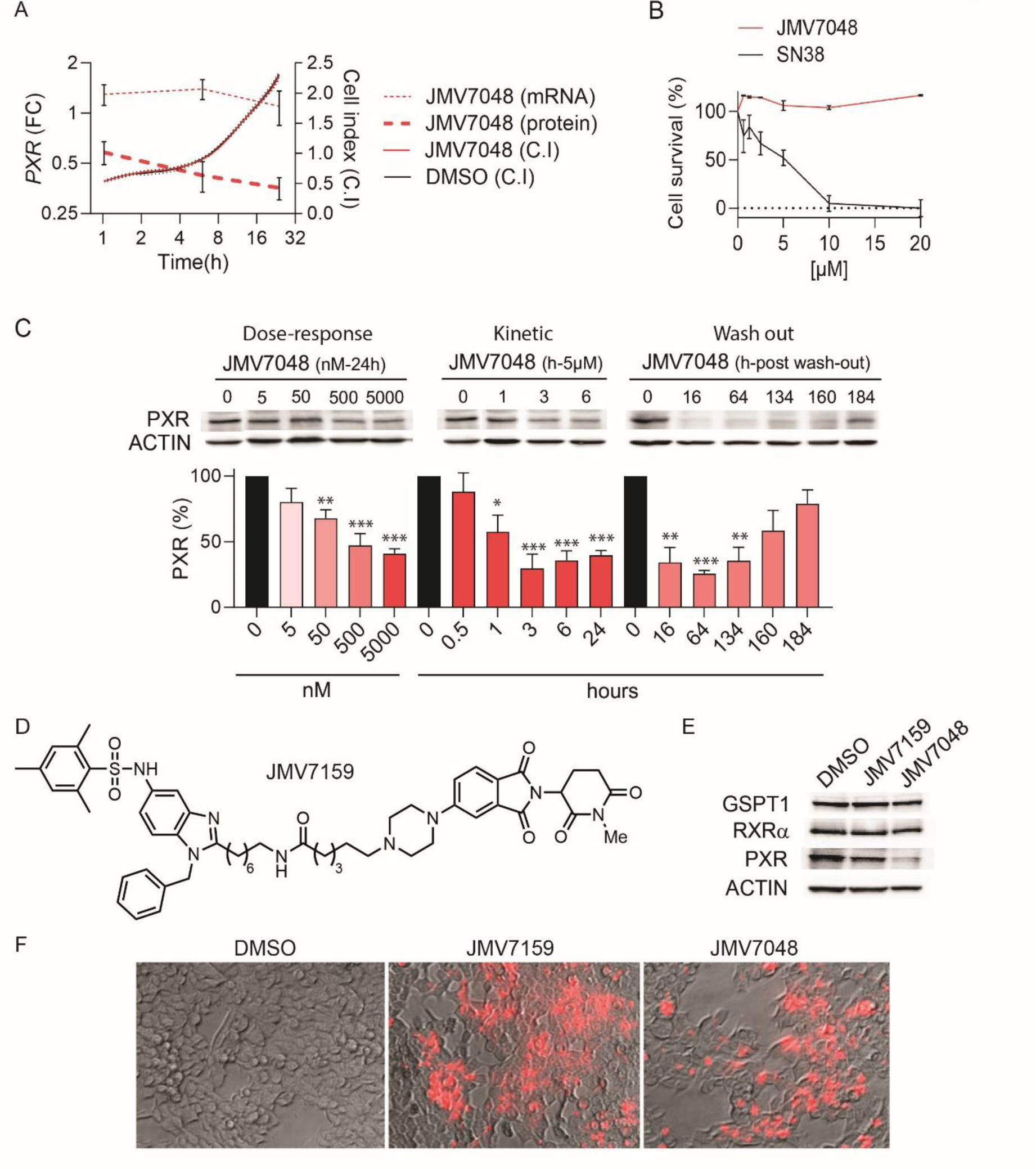
JMV7048 is a PXR degrader. **A)** Parallel analysis of LS174T cells proliferation index (C.I., xCELLigence apparatus) and quantification of *PXR* mRNA and protein expression levels after 24 hours treatment with 5µM JMV7048. **B)** Cell viability analysis of LS174T cells treated for 72 hours with increasing concentrations of JMV7048 or SN38. Data are expressed as mean ± SEM (n=3) and normalized to DMSO (%). **C)** Western-blot analysis and quantification of PXR protein expression in LS174T treated according to the indicated JMV7048 concentrations (Dose-response), or at 5μM JMV7048 for different duration indicated in hours (kinetic), or after an initial treatment of 6 hours with 500nM JMV7048 following by its removal and wash-out for different times (wash-out). **D)** JMV7159 structure **E)** Western-blot analysis of PXR, RXR*alpha* (RXRα) and GSPT1 protein expression in LS174T cells treated 24 hours with 5µM JMV7048 or JMV7159. **F)** *In cellulo* detection of JMV7048 and JMV7159 compounds by fluorescence live cell imaging. ***, p<0.0005, **, p<0.005, *, p<0.05.

We then performed a focused characterization of JMV7048-mediated PXR degradation. Firstly, dose dependent and kinetics assays were performed, as depicted in Fig.2C. Data demonstrate that JMV7048 potently and efficiently reduced endogenous PXR protein in LS174T cells, with a half maximal degradation concentration (DC_50_) of 379 ± 12 nM, a maximum degradation efficacy (*D*_Max_) of 62 ± 10% and a degradation time for 50% (DT_50_) of 62 minutes. After subjecting LS174T cells to a 6-hour treatment with 0.5μM JMV7048, a washout experiment revealed a 50% recovery of PXR levels only after 160 hours. Complete reappearance of PXR was observed only 184 hours post-treatment. These results confirm a reversible and long lasting PXR degradation, potentially indicating a PROTAC-mediated catalytic effect of JMV7048. To verify that the PXR degradation is dependent on E3 CRBN, we synthesized the compound JMV7159 (Fig2D), a JMV7048 analog that contains an N-methylation onto the thalidomide moiety preventing its binding to E3 CRBN (Yang *et al*, 2020). At the concentration where the active JMV7048 PROTAC achieved the maximum reduction in PXR expression (i.e. 5mM), JMV7159 had no effect on PXR expression (Fig.2E), demonstrating that the E3 ligand moiety is crucial for successful JMV7048-mediated PXR degradation. Importantly, JMV7048 reduces PXR but not GSPT1 protein (Fig.2E), in contrast to the PXR-degrader reported in the literature^8^. The lack of PXR degradation observed with JMV7159 is not due to defective cell penetration capacity, as both molecules are detected in the cytosol of live cells thanks to their fluorescent properties (Fig. 2F).

Due to the intrinsic fluorescence property of the thalidomide moiety (which interferes with TR-FRET readouts), preventing the determination of the affinity of JMV7048 for PXR or CRBN, competitive experiments were undertaken to confirm the involvement of both PXR and CRBN recruitment in the JMV7048-mediated degradation of PXR. For this purpose, LS174T cells were pre-treated with an excess of the PXR ligand (JMV6944, Fig.3A) or an excess of the E3 CRBN ligand JMV6945 (Fig.3B). As shown in Fig.3C both competitors entirely inhibited the PXR degradation induced by JMV7048. Secondly, we used E3 ligase and 26S proteasome inhibitors to confirm the involvement of the ubiquitin proteasome machinery in JMV7048-mediated PXR degradation (Fig.3D). Cotreatment of LS174T cells with MLN4924 (Wu *et al*, 2018) (Fig.3E) or with the Bortezomib (Bonvini *et al*, 2007) (Fig.3F) further demonstrated that the degradation of PXR protein was dependent on both the E3 CRBN ubiquitin ligase and the 26S proteasome machinery respectively. Moreover, PXR immunoprecipitation showed that JMV7048 enhanced bortezomib-mediated accumulation of polyubiquitinated PXR (Fig.3F). Therefore, these results unequivocally show that JMV7048 induces PXR degradation through polyubiquitination of PXR via cereblon and proteasome-dependent pathways.

**Figure 3:**
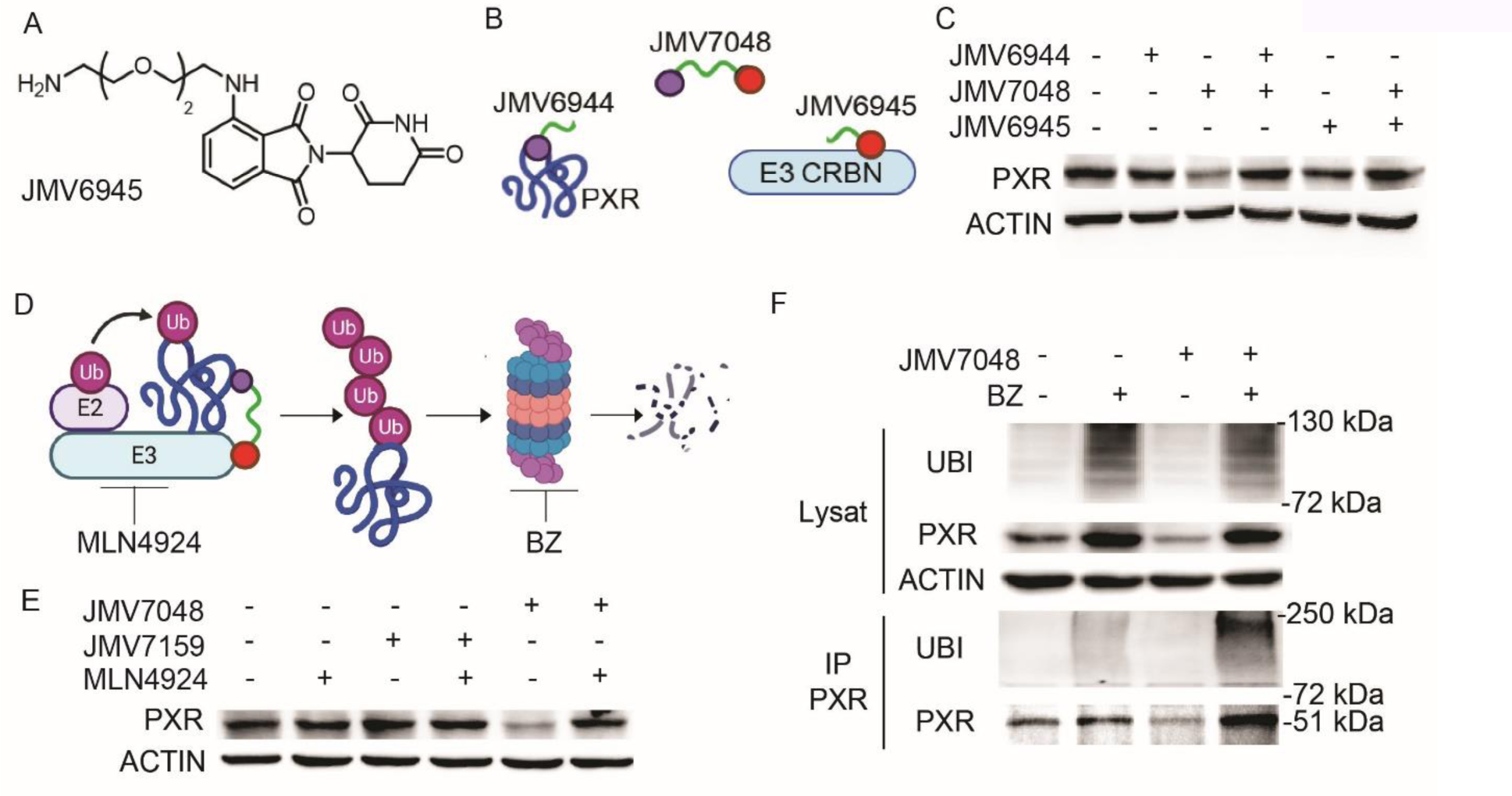
JMV7048 is a bona fide PXR PROTAC. **A)** JMV6945 (CRBN ligand) structure. **B)** Schematic representation of the competitive assays performed in (C). **C)** Western-blot analysis of PXR expression in LS174T cells treated for 24h with 5µM JMV7048 with or without 5µM JMV6944 or JMV6945. **D)** Schematic representation showing the different actors in involved in the proteasome-mediated PXR degradation and their inhibitors (MLN4924 and Bortezomib, BZ) **E)** Western-blot analysis of PXR expression in LS174T cells treated 24 hours with 5µM JMV7048 or JMV7159 with or without CRBN E3 ligase inhibitor (0.5µM MLN4924). **F)** Western-blot analysis of PXR and ubiquitin expression in LS174T cells lysates before and after PXR-immunoprecipitation (IP PXR). Cells were treated for 24 hours with 5µM JMV7048 in presence or absence 100nM Bortezomib (BZ).

### JMV7048 degrades PXR in cancer cell lines but not in primary hepatocytes

We next evaluated the ability of JMV7048 to degrade endogenously expressed PXR protein in several cancer cell lines as well as in primary human hepatocyte cultures. As shown in Fig.4A and Fig.4C, 24 hours of treatment with 5µM JMV7048 significantly reduced PXR expression, not only in LS174T cells (human colon adenocarcinoma), but also in HepG2 (human hepatoma) and AsPC-1 (human metastatic pancreatic adenocarcinoma), two cell lines known to express endogenous PXR (Yokobori *et al*, 2017; Oladimeji *et al*, 2019). In contrast, JMV7048 failed to degrade PXR in primary cultures of human hepatocytes (Fig.4B and 4C). To decipher why JMV7048 cannot degrade PXR in human hepatocytes, we investigated it’s *in vitro* metabolic stability. Accordingly, 400,000 cells (freshly isolated human hepatocytes, HepG2 or LS174T) were incubated with 1µM JMV7048 for five time-points (0, 5, 10, 60 minutes, and 24 hours) and then analyzed by LC-MS/MS. The same experimental protocol was used for all cell lines, allowing a better calculation of both intrinsic clearance values (CL_int_) and half-life (T_1/2_) of JMV7048 in these different models. As shown in Fig.4D, and in agreement with the absence of JMV7048-triggered PXR degradation in human hepatocytes, there was significant acceleration of JMV7048 clearance within hepatocytes compared to colorectal cancer or hepatoma cell lines (8.7- and 5.1-fold respectively). These results suggest a promising heightened vulnerability of JMV7048 to metabolic processes within human hepatocytes (T_1/2_=10h) in contrast to cancer cell lines (with T_1/2_= 3.76 days in LS174T cells and T_1/2_=2.2 days in HepG2 cells). This is in line with the well-known notion that human hepatocytes harbor enhanced levels of drug transporters and drug-metabolizing enzymes compared to cancer cells (Fig.4E).

**Figure 4:**
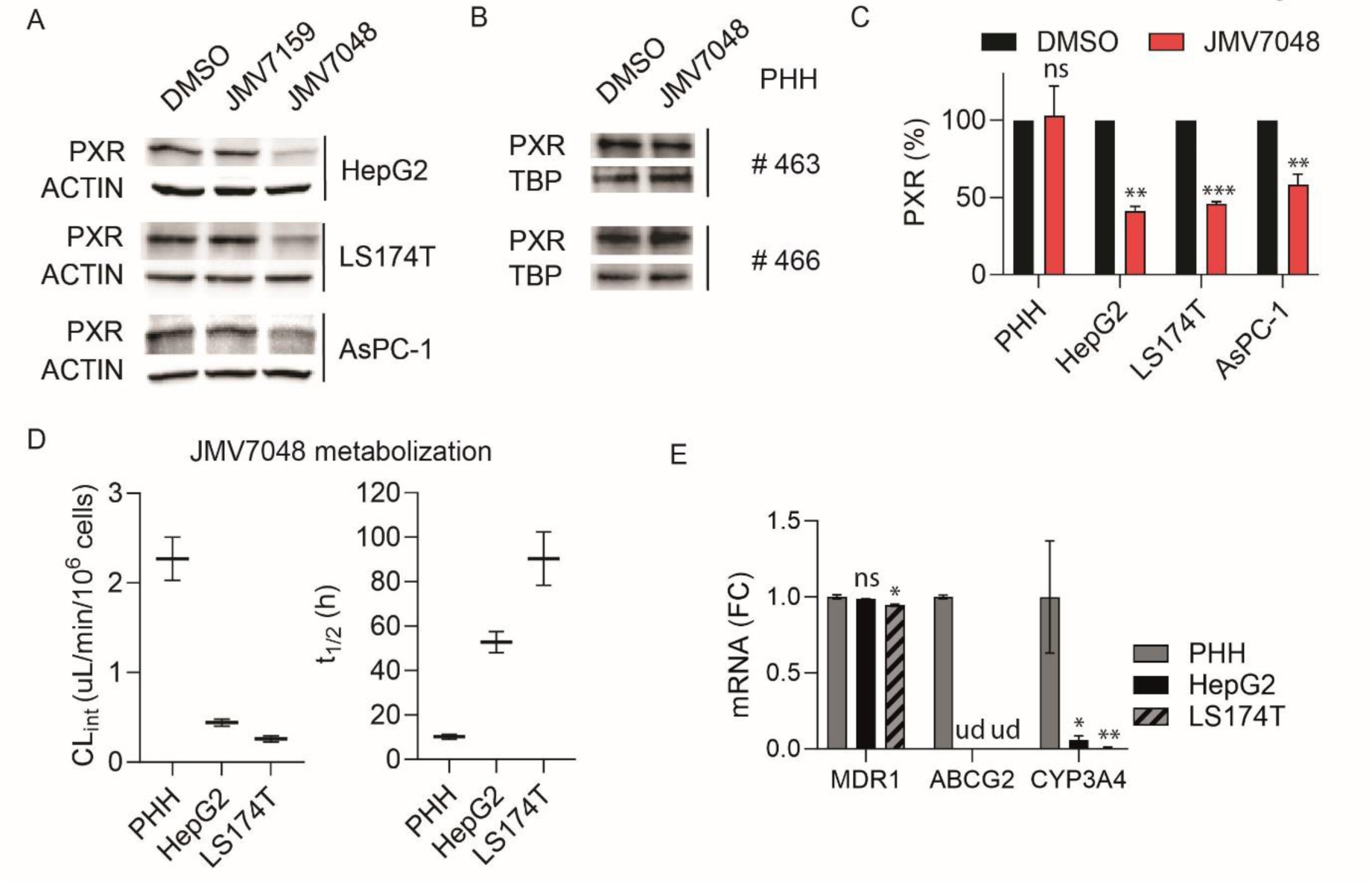
JMV7048 degrades PXR in pancreas and hepatoma cell lines but not in human hepatocytes. **A)** Western-blot analysis of PXR expression after 24 hours treatment with 5 µM JMV7159 or JMV7048 in LS174T, ASPC-1 or HepG2 cell lines. **B)** Western-blot analysis of PXR expression after 24 hours treatment with 5 µM JMV7048 in Primary cultures of Human Hepatocyte (PHH). **C)** Western-blot quantifications of (panels A and B) data are expressed as mean ± SEM (n ≥3) of PXR/ACTIN ratio and normalized to DMSO (%). **D)** Metabolic analysis (intrinsic clearance values -CL_int_- and half-life -T_1/2_-) of JMV7048 in Human Hepatocyte (PHH), LS174T or HepG2 cells treated with 1µM JMV7048 for 5min, 10 min, 1h or 24h (n=2). **E)** RT-qPCR analyses of *MDR1, ABCG2, and CYP3A4* mRNA expression in Human Hepatocyte (PHH), HepG2 and LS174T cells (F.C., Fold Change). Data are expressed as mean ± SEM (n=3) and normalized to PHH condition. ***, p<0.0005, **, p<0.005, *, p<0.05.

### JMV7048 reduces the number of colon cancer stem cells

As cancer stem cells (CSCs) are known to express more PXR and drug-metabolizing enzymes than non-CSC (Planque *et al*, 2016), we then tested the efficacy of JMV7048 in the ALDH-positive CSC subpopulation. Indeed, the high ALDH enzymatic activity (ALDH^+^) (Giraud *et al*, 2016) population is enriched in CSCs. LS174T cells were first labeled with an Aldefluor kit (STEMCELL Technologies) and ALDH-negative and ALDH-positive cells were purified by flow cytometry and then treated for 24 hours with 5µM JMV7048. Western blotting analysis confirmed the overexpression of PXR in CSCs and revealed that JMV7048 induces its degradation in ALDH-positive cells Fig.5A. Having demonstrated that JMV7048 promotes PXR degradation in CSCs, we then investigated its effect on CSC population properties such as ALDH activity, self-renewal and tumor initiation. For this purpose, we treated patient-derived colorectal cancer cells established in our lab, either from primary (CPP1, CPP14) or metastatic (CPP19) tumors, with 5µM JMV7048 for 48 hours. As shown in Fig.5B, the proportion of cells exhibiting high ALDH activity was significantly decreased in all tested cells. Accordingly, treatment with JMV7048 diminished their ability to form spheres in non-adherent conditions (typically used as a readout for challenging the self-renewal capability of CSCs (Hu & Smyth, 2009)), whereas its negative analog (JMV7159) did not yield the same effect (Fig.5C). To continue the impact assessment of PXR degradation on CSC properties, we then tested JMV7048-treated cells for tumor initiation in subcutaneous xenografts, recognized as the gold standard assay. CPP1 cells were first treated *in vitro* for 48 hours with 5µM of JMV7048 before being dissociated. Viable cells (SYTOX-blue negative cells) were purified by flow cytometry and then injected into nude mice (500 or 5000 cells/mouse, n=5/group) for *in vivo* tumor initiation assays (den Hollander *et al*, 2022). As shown in Fig.5D, JMV7048 dramatically reduced the number of mice developing tumors. The calculated tumorigenicity (frequency of tumor-initiating cells) of colon cancer cells was significantly reduced by JMV7048 (1/3557) compared to DMSO-treated cell (1/311, p=0.001).

**Figure 5:**
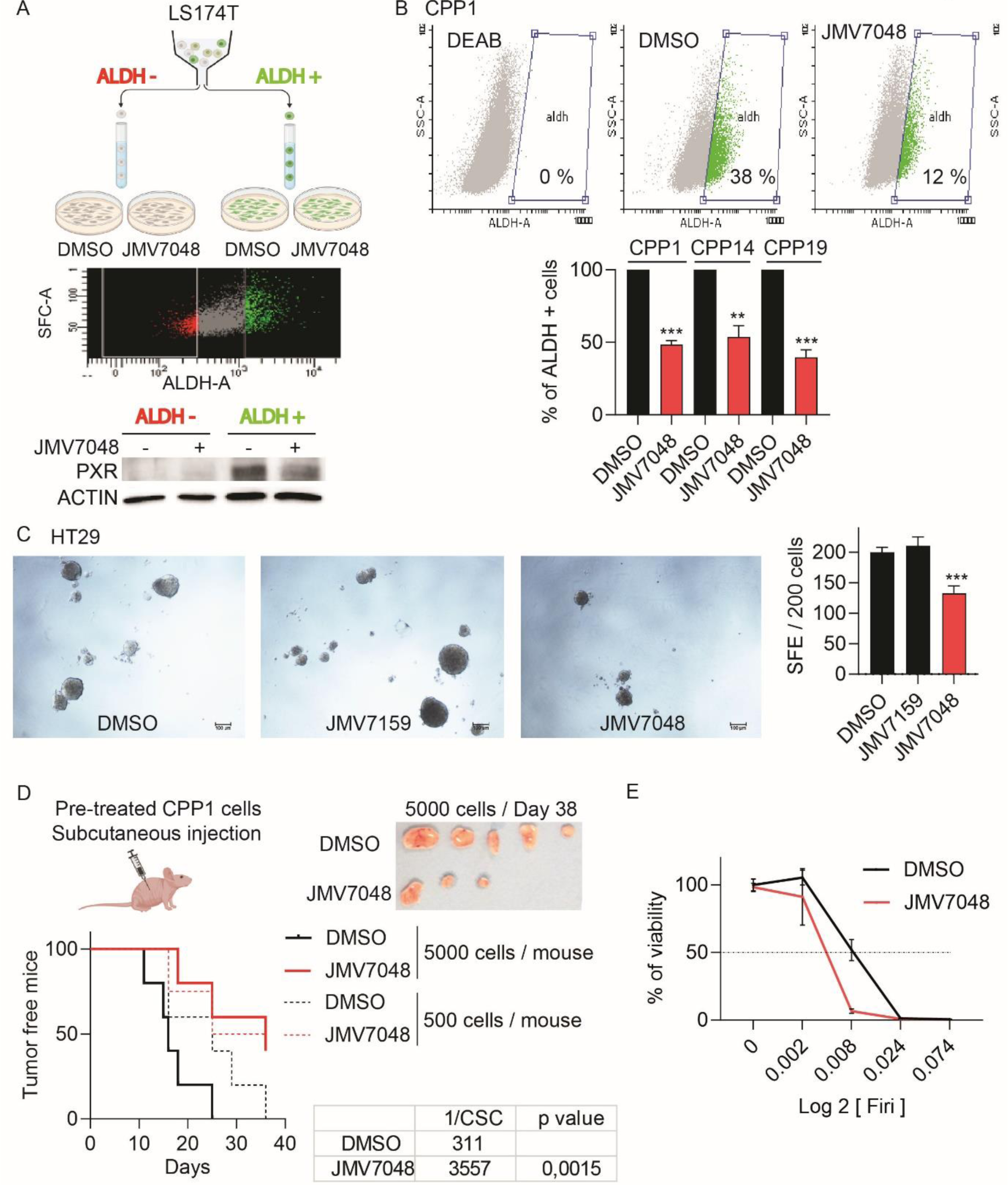
JMV7048 decreases PXR expression in colon CSCs and inhibits CSC population *in vitro.* **A)** Western blot analysis of PXR expression in *Aldefluor*-positive and negative sorted cells and then treated for 24h hours with 5µM JMV7048. **B)** Quantification of *Aldefluor-*positive cells in CPP1, CPP14 and CPP19 patient derived CCR cells after 24 hours treatment with 5µM JMV7048. **C)** Sphere-Forming Efficiency (SFC) assay of HT29 cell line incubated with 5µM JMV7159 or JMV7048. **D)** Log-rank Mantel-Cox Tumor initiation test. CPP1 cells were first treated *in vitro* for 48 hours with 5μM JMV70048 and were injected in Nude mice (500 or 5000 cells/mouse, n=5). **E)** Cell viability analysis of HT29 spheroids pre-treated 24 hours with 5μM JMV7048 and then co-treated with Firi at 0.008X for HT29 and 1.25X for CPP1 (1X = 50µM 5FU + 5µM SN38, n=3). ***, p<0.0005, **, p<0.005.

Because PXR drives the expression of a large network of genes instrumental CSC chemoresistance (Planque *et al*, 2016; Bansard *et al*, 2022), we then assessed the impact of JMV7048 on the survival of colorectal cancer cells treated with chemotherapy *in vitro*. HT29 cells were maintained as spheroids (Kanwar *et al*, 2010) in order to increase the pool of CSCs and thus PXR expression. Cells were first pre-treated for 48h with 5µM JMV7048 and then challenged with increasing concentrations of a combination of 5-FU and SN38 (hereafter named “Firi”, 1X= 50μM 5-FU + 5μM SN38,) as SN38 is the active metabolite of irinotecan. Cells were then incubated for three days before cell viability measurement. As shown in Fig.5E, JMV7048 pre-treatment decreased HT29 (EC_50_DMSO= 0.008X and EC_50_JMV7048= 0.003X, p=0.001) survival after 72h of treatment with Firi. These results demonstrate that, JMV7048 can target CSCs, leading to a decrease in PXR expression, as well as alterations in their self-renewal and chemoresistance.

### JMV7048 inhibits PXR expression and tumor recurrence in xenografted tumors

We first conducted pharmacokinetic studies of JMV7048 in mice following 25 mg/kg intravenous (I.V.), 50 mg/kg intraperitoneal (I.P.) and 50 mg/kg oral (P.O.) administrations). As shown in Fig.6A, although a single I.P. or I.V. administration of JMV7048 resulted in satisfactory drug exposure in plasma, leading to favorable Cmax (maximum plasma concentration) and t_1/2_ values, the P.O. delivery method was largely ineffective, demonstrating suboptimal plasma exposures (AUCt = 75 ng/mL*h). We next evaluated the potential toxicity of prolonged I.V. injections of JMV7048 (25 mg/kg/day, 5 days a week for 15 days) in immunodeficient nude and NOD/scid mice. Throughout this study, we closely monitored changes in body weight, a crucial indicator of drug toxicity (Lewis *et al*, 2002). As depicted in Fig.6B, we observed a generally inconsequential fluctuation in body weight within the JMV7048-treated groups compared to the control groups in both strains of mice. Neither did we notice any clinical signs of toxicity (diarrhea, skin ulcers, hyper/hypoactivity, or changes in motor activity), suggesting that JMV7048 is well tolerated by these mice.

**Figure 6:**
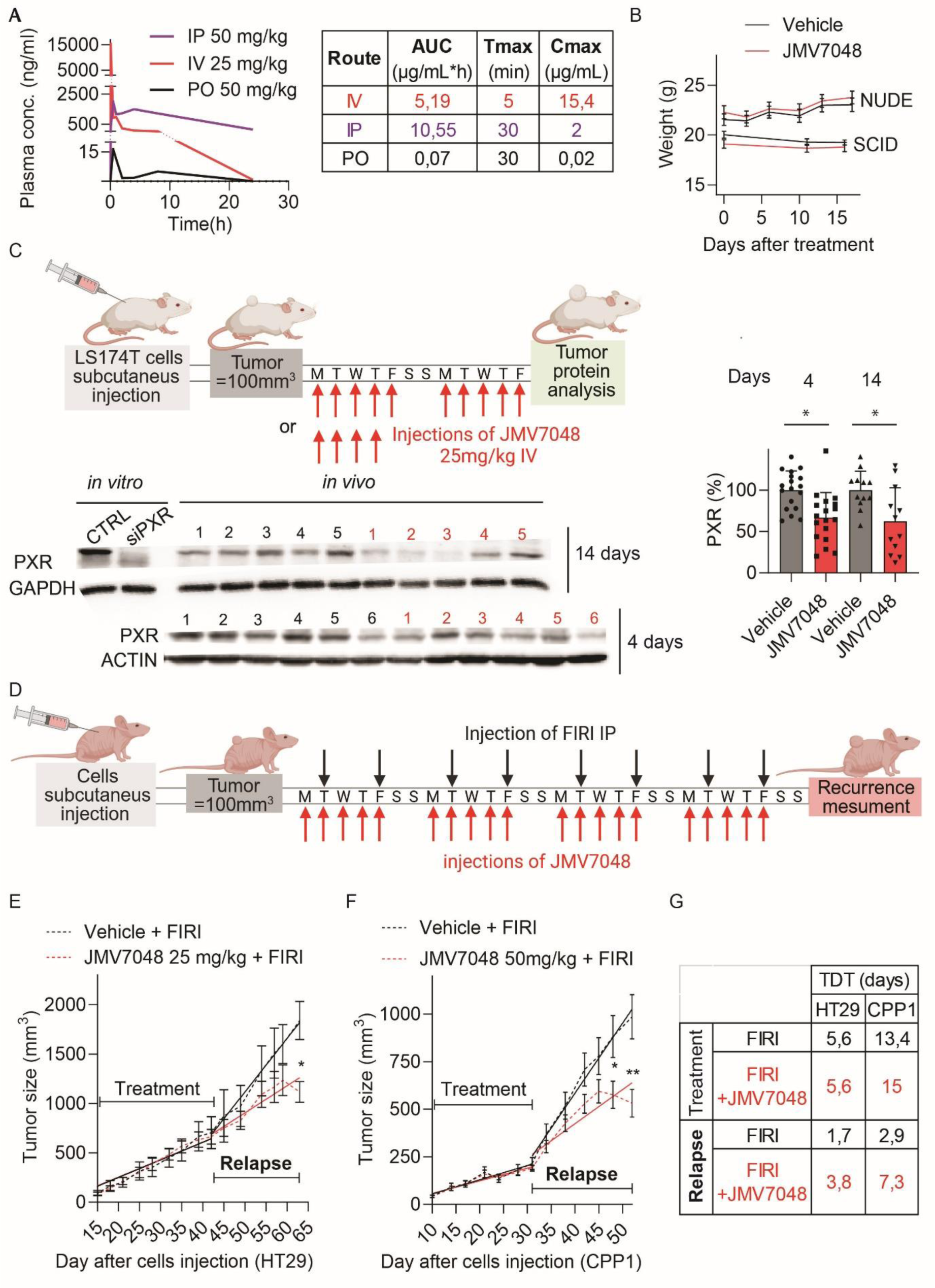
JMV7048 decreases PXR expression and tumors relapse *in vivo.* **A)** Pharmacokinetic studies (plasma concentration profiles, area under the plasma concentration versus time -AUCt-, maximum plasma concentration -Cmax-, and time to reach Cmax -Tmax) of JMV7048 in mice after intravenous (IV), intraperitoneal (IP) or oral (PO) administration. **B)** Body weight analysis of nude and NOD/scid mice treated with or without JMV7048 (25 mg/kg, I.V., 5 times a week for 2 weeks). **C)** Upper panel: schematic representation of *in vivo* treatment protocol: after subcutaneous xenograft in Scid mice of 300.000 LS174T cells, mice were treated with or without JMV7048 (25 mg/kg, I.V., for 4 or 14 days) when tumor volume reached 100mm^3^. Lower panel: Western blot analysis and quantification of PXR protein in resected xenograft tumors (n=10 or 4 /condition 4 days and 14 days respectively). Data (PXR %) represent human-PXR/human-ACTIN ratio and normalized to vehicle treatment. **D)** Schematic representation of *in vivo* treatment protocol: after subcutaneous xenograft of 20.000 of cells (from HT29 or CPP1 spheroids) in athymic nude mice, when tumor volume reached 100mm^3^, mice where treated with FIRI (50mg/kg 5-FU and 25mg/kg irinotecan, IP, twice a week) with or without JMV7048 (IV or IP, 5 days a week) during 4 or 3 weeks. **E)** Tumor volume analysis of HT29 xenograft in mice treated during 4 weeks with FIRI and JMV7048 (25/mg/kg, IV) and after treatment cessation (Relapse). **F)** Tumor volume analysis of CPP1 xenograft in mice treated during 3 weeks with FIRI and JMV7048 (2X25mg/kg/day, IP) and after treatment cessation (Relapse). **G)** Calculated tumor volume doubling times (TDT) during and after (Relapse) treatments.

We subsequently explored the efficacy of JMV7048 in preclinical mouse xenograft models. We subcutaneously xenografted 300,000 LS174T cells in NOD/scid mice and allowed tumor volume to reach 100 mm^3^. Mice were then exposed to I.V. JMV7048 treatment (25 mg/kg/day, 5 days/week - i.e. Mondays to Fridays) for 4 days or 2 weeks (Fig.6C). Western blotting analyses were then performed to determine PXR expression in excised xenograft tumors. As depicted in Fig.6D, JMV7048 induced a significant (_∼_50%) PXR degradation after 4 or 14 days of treatment, demonstrating that JMV7048 can be absorbed and distributed into tissues, at least in subcutaneous xenografts of human tumor cells, and retain its degradation activity *in vivo*. We finally tested the impact of JMV7048 in post-treatment relapse using two preclinical mouse xenograft models and using two different administration routes (I.V. *versus* I.P.). For this purpose, we subcutaneously xenografted HT29 (Fig.6E) or CCP1 patient-derived (Fig.6F) cells (20000 cells/mouse). Once tumors had reached the volume of 100 mm^3^, mice were randomized and received the following treatment regimens: FIRI alone (50 mg/kg 5-FU and 25 mg/kg irinotecan, twice a week), or combined JMV7048 (25 mg/kg/day I.V. for HT29 cells and 25 mg/kg/twice a day I.P. for CCP1 cells, 5 days/week) *plus* FIRI. In assessing the dynamics of tumor growth, we utilized simple linear progression models to analyze the tumor volume doubling time (VDT) both during and following treatment for each group. As shown in Fig6G, we observed that JMV7048 did not significantly affect tumor growth during FIRI treatment in HT29 or CRC1 xenograft models. For instance, during FIRI treatment, the calculated volume doubling time (VDT) for HT29 xenografts was 5.6 days regardless of the presence or absence of JMV7048. In CCP1 xenografts, the VDT was slightly slower in presence in presence of JMV7048 (15 days *versus* 13.3 days for FIRI treatment alone). Signs of tumor relapse became apparent in both xenograft models shortly after treatments cessation, notably in the control groups (FIRI alone), as indicated by a rapid increase in tumor doubling. For instance, tumor volume doubling time increased by more than threefold for HT29 cells (from 5.6 days to 1.8 days) and 4.6-fold for CCP1 cells (from 13.4 days to 2.9 days) following the cessation of FIRI treatments. Nonetheless, the mice treated with FIRI *plus* JMV7048 demonstrated a delayed tumor relapse in comparison to the mice exclusively treated with FIRI. Notably, the tumor growth subsequent to JMV7048 treatments was nearly 2.1-fold and 2.5-fold slower when compared to the control FIRI groups in HT29 and CCP1 xenograft tumors, respectively. These findings collectively demonstrate that JMV7048 acts as a promising adjuvant therapy combined with chemotherapy, postponing tumor relapse in preclinical models.

## DISCUSSION

Although considerable progress has been made in diagnosis and treatment, colorectal cancer remains mostly insensitive to therapy. Accumulating studies have demonstrated that cancer stem cells (CSCs) are one of the reasons for relapse and metastasis (Prud’homme, 2012; Adorno-Cruz *et al*, 2015). Indeed, because of their innate chemoresistance properties, CSCs can withstand conventional cancer treatments, allowing them to survive and regenerate the tumor, leading to disease recurrence (Kreso & Dick, 2014). However, while targeting CSCs appears promising, only a few anti-CSC molecules have successfully completed phase III trials. Indeed, most anti-CSC strategies developed so far have aimed to block fundamental stemness signaling, with therapeutic limitations due to their impact on normal stem cells (Takebe *et al*, 2015). In previous studies, we demonstrated that relapse-prone CSCs are characterized by high PXR expression and activity. In addition this nuclear receptor drives the expression of a large network of genes that are crucial for CSC chemoresistance (Planque *et al*, 2016; Bansard *et al*, 2022). Thus, we suggested that decreasing PXR expression/activity using pharmacological inhibitors may represent a promising strategy to improve the efficiency of conventional chemotherapy in colorectal cancer patients through the sensitization of CSCs. We then focused on developing druggable molecules that could be used as adjuvant to chemotherapy. However, the development of PXR antagonists or inhibitors is relatively challenging due to the nature of its ligand binding domain. We consequently shifted our focus to a more innovative strategy, the proteolysis targeting chimera (PROTAC).

PROTACs represent an innovative class of compounds that overcome traditional limitations, such as targeting low affinity and low specificity undruggable receptors, opening a new therapeutic modality. But in the same time, they challenge the rules used so far for drug discovery. For instance, their high molecular weight (>1000Da) and the linker instability at the attachment points to ligands, make the delivery and bioavailability of PROTACs the most significant hurdles to overcome on the way to the clinic (Neklesa *et al*, 2017). Accordingly, to date, only two PROTACs have advanced to Phase II clinical trials (https://www.arvinas.com/research-and-development/pipeline/) for the treatment of metastatic castration resistant prostate cancer (Androgen Receptor: NCT03888612) and locally advanced metastatic breast cancer (Estrogen Receptor: NCT04072952). However, the PROTAC technology is still maturing, and the design elements for successful PROTAC-based drugs are constantly evolving. In this context, we innovatively developed a degrader combining a high affinity PXR agonist, that we previously identified (Benod *et al*, 2008), with an E3 ligase ligand. So far, PROTACs were generally constructed from antagonists of the target protein, with very few exceptions (Zhou *et al*, 2022; Xu *et al*, 2021), however PXR antagonists are very rare, and unfortunately characterized by affinities on the order of micromolar (µM). Moreover, the first attempt to create a PXR PROTAC from an antagonist led to the generation of a GSPT1 molecular glue instead of a bona fide PXR PROTAC^8^. Thanks to crystallization and LBD PXR dynamics studies (Delfosse *et al*, 2021), we successfully developed the first PXR PROTAC degrader, JMV7048, which can induce durable degradation of PXR *in vitro* and *in vivo* and exhibit therapeutic potential. Unlike previously identified PXR antagonists, which were only effective at the micromolar range and required a direct and sustained interaction with the PXR protein, JMV7048 remains active down to 400 nanomolar, and exhibits catalytically effects that endure for several days. Therefore, our results demonstrate that agonist-based PROTACs are a promising approach to create PROTACs, even in the absence of an appropriate antagonist.

Our results indicate that JMV7048, in combination with chemotherapy, decreases the pool of chemoresistant CSCs, delaying tumor relapse in mouse xenograft models, providing the proof of concept that PXR PROTAC approach is an innovative adjuvant strategy to sensitize cancer stem cells. Our investigation, we noticed that while JMV7048 significantly decreased PXR protein expression in colon, hepatoma or pancreatic cancer cell lines, it was inefficient in primary culture of human hepatocytes. Accordingly, compared metabolic stability analysis showed an enhanced vulnerability of JMV7048 to metabolic processes within human hepatocytes in contrast to cancer cell lines. It is well known that human hepatocytes contain all hepatic drug-metabolizing enzymes and cofactors at physiological levels and represent the “gold standard” for the study of metabolic stability. Therefore, it is likely that JMV7048 underwent more rapid metabolization in the liver compared to cancer cells. These observations suggest that 1) JMV7048 will not interfere with the crucial functions of PXR in the liver, thereby preserving the vital hepatic drug-metabolizing functions of PXR, and 2) that JMV7048 might be rapidly inactivated thought the first-pass effect. Thus, demonstrating the selectivity of JMV7048 in its action on PXR between cancer cells and normal hepatocytes can contribute significantly to its development as a targeted therapeutic agent, potentially providing a more effective and safe treatment option for several types of cancer without compromising normal liver function and drug metabolism. In the order hand, the enhanced liver first-pass effect may decrease the active JMV7048’s concentration before reaching the systemic circulation and thus it will require increased oral dosages or alternative administration routes, such as parental administration, as employed in our preclinical tests.

Finally, we observed that JMV7048 is well tolerated, even at 50mg/kg/day for 3-4 weeks, and does not produce any clinical signs of toxicity in mice. Despite its relatively poor pharmacokinetic properties, JMV7048 significantly decreased PXR expression and delayed tumor relapse in xenograft tumor models. These observations support previous studies indicating that PROTACs establish a disconnection between pharmacokinetics and effectiveness (Bondeson *et al*, 2015). Indeed, PROTACs have been reported to operate catalytically and do not require complete drug-receptor binding for their effectiveness, often necessitating high drug concentrations and this is particularly beneficial for low-affinity receptors such as PXR. There is now the need for extensive efforts to improve the *in vivo* efficiency and stability of this molecule but we maintain the belief that the development of a PXR PROTAC provides a novel avenue to potentiate cancer therapy and address critical challenges associated with tumor relapse in colon cancer and other types of cancer, where PXR is expressed and associated with poor clinical outcomes. In addition, the development of PXR PROTACs could be explored in diseases or metabolic disorders known to be associated with dysregulation of PXR activity such as such non-alcoholic fatty liver disease (NAFLD) or type 2 diabetes.

## METHODS

### PXR crystal structures

The human PXR-LBD was produced and purified as described in bioRxiv (in preparation). Briefly, the coding region of the PXR-LBD (130-434) was expressed in E. coli as a fusion protein containing a fragment of the steroid receptor coactivator-1 (SRC-1, 678-700) in its C-terminal part to improve PXR stability. After an initial purification step by Ni-affinity chromatography, the SRC-1 fusion moiety was cleaved thanks to a thrombin cleavage site prior to size exclusion chromatography. The purified protein was then concentrated at 4 mg.ml^-1^. Crystals of PXR complexes with JMV6845 and with JMV6944 were obtained by co-crystallization. 1 μl of protein with 3 molar equivalents of ligand was mixed with 1 μl of precipitant (100 mM imidazole pH 7.0-7.4, 10-13% (v/v) isopropanol), and let equilibrated against a reservoir of 500 μl of precipitant. Crystals appear in 24h. Diffraction data were collected on the ID30A-3 (JMV6845) and ID23-2 (JMV6944) beamlines at the European Synchrotron Radiation Facilities (λ = 0.96769 Å and λ = 0.87313 Å, respectively, 100K), Grenoble, France. Data were processed and scaled with XDS and XSCALE (Kabsch, 2010). Crystals belong to space group P 43212. The structure was solved and refined using Phenix (Afonine *et al*, 2012) and COOT (Emsley *et al*, 2010). The percentage of residues located in the favored Ramachandran plot region is 98.9% and 99.2% for the JMV6845 and the JMV6944 complexes respectively (calculated with MolProbity (Williams *et al*, 2018)). Data collection and refinement statistics are summarized in Supplementary Table 1. Figures were prepared with PyMOL (http://pymol.org/).

### TR-FRET PXR Competitive Binding Assay

TR-FRET PXR (SXR) Competitive Binding Assay Kit (LanthaScreen™) was used to analyze the binding of PXR agonists to the LBD of PXR accordingly to the manufacturer’s instructions. Briefly, reaction mixture was incubated for 2 hours at room temperature in a 384-well microplate. Fluorescence reading was carried out using a PHERAstar (BMG Labtech).

### Cell lines and patient-derived cells culture

The following cell lines were obtained from the ATCC: LS174T (human colon adenocarcinoma, CL-188), HT29 (human colon adenocarcinoma, HTB-38), AsPC-1 (human pancreas Adenocarcinoma, CRL-1682), HepG2 (human liver carcinoma, HB-8065).

The following cell lines were obtained from CCR patient biopsies: CPP1 (human colon adenocarcinoma), CPP14 (human colon adenocarcinoma), CPP19 (metastatic human colon adenocarcinoma). These biological samples were provided by CHU-Carémeau (Nîmes, France, ClinicalTrial.gov Identifier#NCT01577511) and patients’ clinical characteristics are presented in Supplementary Table 2.

Primary human hepatocytes (PHHs) were isolated as described previously (Pichard *et al*, 2006) from donor organs unsuitable for transplantation or from liver resections performed in adult patients for medical reasons unrelated to our research program. Liver samples were obtained from the Biological Resource Center of Montpellier University Hospital (CRB-CHUM; http://www.chu-montpellier.fr; Biobank ID: BB-0033-00031) and this study benefitted from the expertise of Dr Benjamin Rivière (hepatogastroenterology sample collection) and Dr Edouard Tuaillon (CRB-CHUM manager). The patients’ clinical characteristics are presented in Supplementary Table 3.

LS174T and AsPC-1 cell lines were cultured in RPMI (Thermo Fisher Scientific) containing 10% FBS (Eurobio). HT29, CRC1, CPP19 cell lines were cultured in DMEM (Thermo Fisher Scientific) containing 10% FBS. The HepG2 cell line was cultured in MEM (Life) containing 10% FBS. All cell cultures were incubated in a humidified atmosphere with 5% CO2 at 37C and divided once reaching 80% confluency. Low passage numbers’ (<12 generations) cells were used. Cell lines were verified to be free of mycoplasma and human (hIMPACT Profile III) pathogens by PCR testing (Idexx Bioanalytics). Hepatocytes were seeded in collagen-coated dishes at 2.1x 10^5^ cells/cm² in ISOM medium containing 2% bovine serum (Isom et al., 1985). Following 24 hours of attachment medium was changed to ISOM without serum. For spheroid cultivation, the cells are seeded in flasks or wells coated with poly-HEMA (Sigma) at low density (1 cell/µl) in DMEM/F12 medium (Thermo Fisher Scientific) supplemented with Supplement N2 (Thermo Fisher Scientific), hEGF (Miltenyi), hFGF (Miltenyi), glucose 0.3% and insulin 0.002% (Sigma).

For the cultivation of tumoroids, the cells are seeded in 96-well plates at 1000 cells in 50ul DMEM/matrigel (v:v) (Corning,) then incubated at 37°C for 20 min then the domes formed are covered with 100μl of DMEM + Glutamine + 10% FBS medium.

### Cell viability assays

Cells were plated at 2,000 cells per well in 96-well plates in DMEM or RPMI with 10% FBS. After 24 hours, cells were treated with compounds for 72 hours. Cell viability was assessed by Sulforhodamine B staining as previously described (Bansard *et al*, 2022).

### Proliferation assays

The xCELLigence system makes it possible to measure the effect in real time toxicity of a molecule on cell proliferation. LS174T cells were plated at 2,000 cells per well in 8-well microplates in RPMI with 10% FBS and treated with JMV7048 or DMSO for 72 hours. Cell proliferation was assessed by cell index calculation.

### Transfection assay and oligonucleotides

PXR siRNA (siRNA NR1I2 silencer human, s16909, Life) transfection experiments were performed using Lipofectamine RNAiMax (Invitrogen) according to the manufacturer’s instructions.

### Luciferase activity

The LS174T cells stably expressing the PXR coding region expression cassette, and both CYP3A4 promoter-luciferase and CMV-driven eGFP constructs have been described previously (Planque *et al*, 2016). Cell lines were cultured in RPMI plus 5% FBS Treated with Activated Carbon (Eurobio). Cells are lysed with Passive Lysis Buffer (Promega) and GFP is read for viability data in a Tecan apparatus, then the luciferase substrate is added to read the luminescent signal (luc). Data are expressed as a ratio luc/eGFP.

### RNA isolation and RT-PCR assays

#### Cells lines

Total RNA was extracted using the RNeasy mini kit (Qiagen) and treated with DNAse-1 as recommended by the manufacturer. The first strand cDNA was synthesized using Superscript II (Invitrogen) and random hexamers, and gene expression was measured by real-time PCR.

#### Primary Human Hepatocyte

After extraction with Trizol reagent (Life Technologies), 500 ng of total RNA was reverse-transcribed using random hexamer and MMLV Reverse Transcriptase Kit (Life Technologies).

Quantitative polymerase chain reactions were performed using the Roche SYBER Green reagent and a LightCycler 480 apparatus (Roche Diagnostic, Meylan, France). Each cDNA sample was amplified in triplicate using SYBR Green (Roche) on the LC480 real-time PCR system (Roche). Primer sequences are presented in Supplementary Table 4. The following program was used: one step at 95°C for 10 min and then 50 cycles of denaturation at 95°C for 10 s, annealing at 62°C for 15 s and elongation at 72°C for 15 s.

### Protein analysis

Cell lines: Cells were lysed with RIPA buffer and prepared for total protein extraction with protease inhibitors (Roche). After 20 minutes of centrifugation (>16000g), the supernatant was collected and protein concentration were determined by the bicinchoninic acid method (Pierce Chemical, Rockford, IL). Bovine serum albumin (Pierce Chemical) was used as standard. Ninety µg of protein were subjected to 10% SDS-PAGE and transferred to nitrocellulose membrane (Amersham). The following antibodies were used: GADPH (sc-32233, final dilution: 1/5000), RXR-alpha (sc-515929, final dilution: 1/500), CYP3A4 (sc-53850, final dilution: 1/500), Ubiquitin (sc-166553, final dilution: 1/500) and PXR (sc-48340, final dilution: 1/500) from Santa Cruz, GSPT1 (ab126090, final dilution: 1/500) and actin (ab253283, final dilution: 1/1000) from abcam and Actin (A4700, final dilution: 1/5000) from sigma. Bands intensities were measured on an image with Image Lab software (BIORAD).

Primary Human Hepatocyte: Cytosolic and nuclear fractions were prepared using the NE-PER cell fractionation kit (Thermo Fisher) according to the manufacturer’s instruction. The protein concentration was determined by the bicinchoninic acid method (Pierce Chemical, Rockford, IL). Bovine serum albumin (Pierce Chemical) was used as standard. Twenty µg of nuclear proteins were separated on precast 4-15% SDS-polyacrylamide gels (Bio-Rad Laboratories), and then transferred onto polyvinylidene fluoride membranes (Bio-Rad Laboratories). Membranes were incubated with mouse monoclonal anti-PXR (Santa-Cruz, clone H11) or rabbit anti-TATA-box binding protein (TBP, NB-22-7526, Neo Biotech, Clinisciences, France) antibodies. Immunocomplexes were detected with horseradish peroxidase-conjugated rabbit or mouse secondary antibodies (Sigma) followed by enhanced chemiluminescence reaction (Millipore, Molsheim, France). Chemiluminescence was monitored using a ChemiDoc-XRS^+^ apparatus (Bio-Rad Laboratories) and quantified using Image Lab software (version 6.1).

Xenografts tumors: snap frozen tumor pieces (±300mm^3^) were placed in a tube containing ceramic beads (Matrix D, MP Biomedicals) with 400μl of RIPA. The samples were subjected to 4 cycles of lyses in FastPrep-24^TM^ apparatus (MP Biomedicals) for 20 sec followed by incubation for 5 min on ice. After 20 minutes of centrifugation (>16000g), the supernatant was collected and protein concentration was determined by the bicinchoninic acid method (Pierce Chemical, Rockford, IL). Bovine serum albumin (Pierce Chemical) was used as standard. Ninety µg of protein were subjected to 10% SDS-PAGE and transferred to nitrocellulose membrane (Amersham).

### Immunoprecipitation

LS174 PXR cells were plated at 4.10^6^ cells in 175mm^2^ flask in RPMI with 10% FBS. After 6 hours, cells were treated with compounds for 16 hours. Cells were lysed with 400uL RIPA buffer and prepared for total protein extraction with protease inhibitors (Roche). One mg of protein was precleared with A/G agarose beads for 30 minutes before PXR antibody (1.6μg) incubation (sc-48340, Santa Cruz) for 24h. Immunoprecipitation was performed with A/G agarose beads for 4h, beads were then extensively washed 3 times with TBS Tween 0.01% and 1 time with RIPA before resuspension in 50μL 2X Laemly Buffer.

### *Aldefluor* assay and fluorescence-activated cell sorting (FACS)

The Aldefluor assay (Stem Cell Technologies) was performed according to the manufacturer’s instructions. ALDH-positive cells and ALDH-negative cells were identified by comparing the same sample with and without the ALDH inhibitor diethylaminobenzaldehyde (DEAB). The gating strategy for flow cytometry analysis of Aldefluor-stained samples was as follows: cells were first stained using the ALDH assay, then stained with SYTOX Blue Dead Cell Stain (Invitrogen). All samples were analyzed by sequential gating including the main population (SSC vs FSC), single cells (SSC-A vs SSC-W), and viable (SYTOX Blue-negative) cells (SSC-A vs SYTOX Blue channel). Dead cells were excluded based on light scatter characteristics and SYTOX Blue Dead Cell Staining. Cells that had been incubated with the DEAB inhibitor were used to set the negative control gate, upon which identification of the ALDH-positive subpopulation in the test samples (without DEAB inhibitor) was based. Cells were sorted using a FACSAria II (BD) and analyzed using Flowing software (v 2.5.1; http://flowingsoftware.btk.fi/). ALDH activity was also analyzed using the MACSQUANT (Miltenyi) analyzer with the protocol defined above.

### Sphere formation assays

Percentage of Cell Forming Spheres was determined after plating 200 cells/well in M11 medium in 96 well plates in ultra-low attachment plates (PolyHeme coating). Spheres were counted between 7 and 10 days if their diameter exceeded 50μM.

### *In vivo* experiments

All experiments were performed according to the European Union (Council directive 86/609EEC) and institutional/local guidelines on laboratory animal usage. Animal protocols were approved by the French ethical committee for animal testing (authorization referral #38102). All efforts were directed at minimizing animal discomfort and to reduce the number of animals used (3R rule). Rodent housing conditions used in this study are: temperature set point: 22°C; high limit: 23°C; low limit: 21°C Humidity set point: 45%; high limit: 55%; low limit: 40%. Light cycle: 12 hours light/dark. No mouse exhibited severe loss of body weight (>15%) or evidence of infections or wounds. Female 4- to 6-week-old athymic nude mice, Crt/NU (NCr)Foxn1nu, (CRC1 and HT-29) BALB/c SCID BCySmn.cb17-prkcdscid/J*) mice (LS174T) were purchased from the Charles River. Cancer cell lines were suspended and counted in cold DMEM or RPMI medium with 50% of matrigel, and injected into mice subcutaneously. Tumor volume ([length x width^2^]/2) was measured with a caliper. Randomization and treatment started once tumor volume reached 100 mm3, and mice were sacrificed when tumors reached 1,500 mm^3^.

#### Tumor initiation assays

CPP1 cells were pre-treated in vitro for 48h at 5uM then viable cells (7AAD negative cells) were resuspended in 100μL cold DMEM medium with 50% of matrigel, and injected into Crt/NU (NCr)Foxn1nu mice subcutaneously.

#### Mice treatment

JMV7048 was dissolved in a vehicle composed by 20 % kolliphor hs 15, 5% EtOH qsp Water with 5% dextrose and injected by codal (IV) or intraperitoneal injections (IP). FIRI cocktail was dissolved in PBS and IP injected into mice at a dose of 50 mg/kg for 5-FU and 25 mg/kg for irinotecan two times a week.

### Statistical analysis

For each experiment, data are shown as mean ± S.E.M of at least three independent experiments. Graphpad Prism7 software was used for data analysis. The Mann Whitney test was used to analyze the difference between two groups of quantitative variables with alpha-value set at 5%.

## Acknowledgements

We acknowledge the contribution of iExplore animal facility (IGF, Montpellier). We thank C. Duperray (IRBM, Montpellier) from the Montpellier RIO Imaging platform for flow cytometry experiments. This work was supported by Grants from the ANR (Agence Nationale de la recherche, France), AxLR (INSERM) and CNRS INSB maturation programs, MITI Prime-80 (CNRS), INCa-Cancéropôle GSO, Association pour la Recherche contre le Cancer (France), Ligue contre le cancer (France), Key Initiative Muse “Biomarkers and Therapy”, GEFLUC and SIRIC of Montpellier (France).

## Author contributions

*G.L, A.C, J.P, M.A and J-M.P. designed the research. L.B. G.L., J.P., M.A. and JM-P, designed part of the experiments. L.B., G.L., V.D., T.H., M.A., E.R., Q.D., S. G-C., M-C., B.L, and A.R.M. performed the experiments. L.B., G.L., V.D., T.H., M.A., E.R., Q.D., S. G-C., M-C., B.L, and A.R.M. analyzed the data. J.P., W.B., M.A., and J-M.P.* wrote the manuscript.

